# Oxygen production by an ammonia-oxidizing archaeon

**DOI:** 10.1101/2021.04.01.436977

**Authors:** Beate Kraft, Nico Jehmlich, Morten Larsen, Laura Bristow, Martin Könneke, Bo Thamdrup, Donald E. Canfield

## Abstract

Ammonia-oxidizing archaea (AOA) are one of the most abundant groups of microbes in the world’s oceans and are key players in the nitrogen cycle. Their energy metabolism, the oxidation of ammonia to nitrite, requires oxygen. Nevertheless, AOA are abundant in environments where oxygen is undetectable. In incubation experiments, where oxygen concentrations were resolved to the nanomolar range, we show that *Nitrosopumilus maritimus* produces oxygen (O_2_) and dinitrogen (N_2_). The pathway is not completely resolved, but it has nitric oxide as a key intermediate. Part of the oxygen produced is directly used for ammonia oxidation, while some accumulates in the surrounding environment. *N. maritimus* joins a small handful of organisms known to produce oxygen in the dark, and based on this ability, we re-evaluate their role in oxygen-depleted marine environments.

## Main Text

Ammonia-oxidizing archaea (AOA) are only known to oxidize ammonia to nitrite using oxygen: NH_3_ + 1.5 O_2_ → NO_2_^-^ + H_2_O + H^+^ (*1, 2*). Yet, AOA are highly abundant in environments with very low or even undetectable oxygen concentrations such as marine oxygen-minimum zones (OMZs) (*3*–*5*). The role of AOA in such environments is enigmatic as they have no known anaerobic metabolism. We used trace luminescence oxygen sensors (hereafter: optodes) (*6*) to explore the physiology of AOA at low nanomolar oxygen concentrations and functional anoxia (oxygen levels below detection) as typically found in OMZs (*7, 8*), discovering oxygen production by the marine AOA *Nitrosopumilus maritimus SCM 1*(*9*). Dark, non-photosynthetic, oxygen production is rare in nature with three known pathways including chlorite dismutation during perchlorate/chlorate respiration (ClO_2_^-^ → Cl^-^ + O_2_), detoxification of reactive oxygen species (e.g. H_2_O_2_ dismutation) and nitric oxide dismutation (2NO_2_^-^ → 2NO → N_2_ + O_2_) (*10*). While not fully resolved, we show that the pathway of oxygen production by *N. maritimus* is none of these and thus novel. Given the abundance of *N. maritimus* in oxygen-sparse environments, anaerobic oxygen production may be common in nature. We also show that oxygen production is linked to N_2_ production and thereby identify a previously unknown and potentially environmentally significant N_2_ production pathway.

We first grew axenic cultures of *N. maritimus* aerobically as an ammonia oxidizer. The cultures were then sparged with argon to oxygen levels below 5μM, where the remaining oxygen was consumed by *N. maritimus* through continued ammonia oxidation, indicating physiologically active cells. Surprisingly, after oxygen was completely consumed, it immediately started to increase again (Fig. 1A). A series of additions of oxygen-saturated water showed the same recurring pattern: oxygen was consumed and increased directly thereafter (Fig. 1A). When no oxygen additions were made, oxygen build-up continued over hours and reached levels of 100-200nM (Fig. 1A). This pattern was observed reproducibly in multiple incubations carried out over 2 years.

**Fig. 1.**
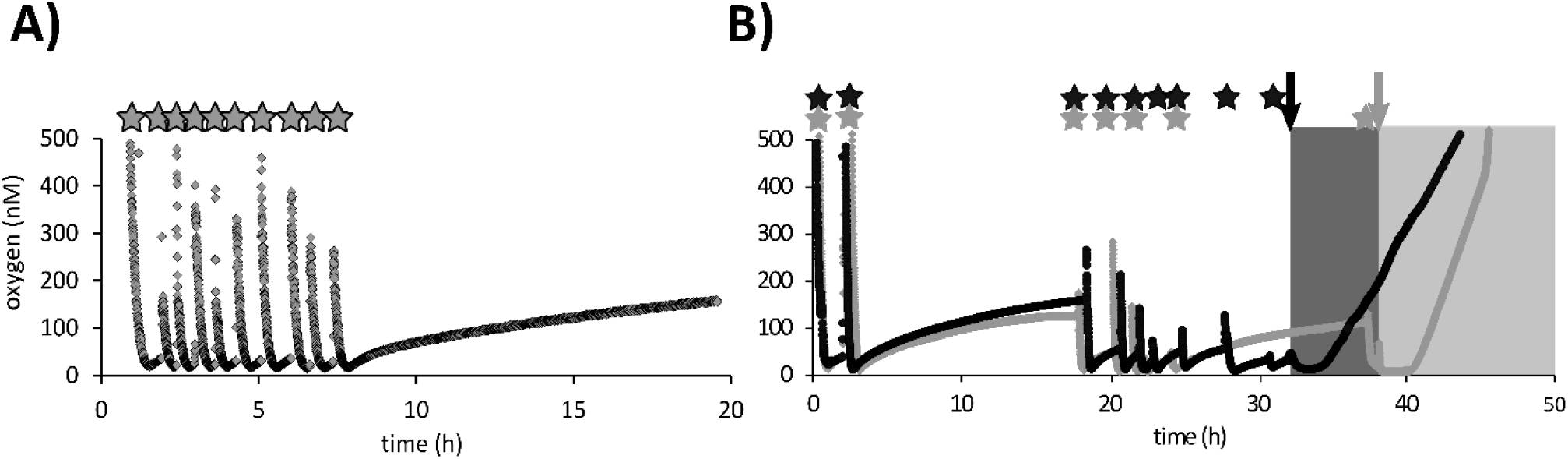
Oxygen production by *N. maritimus*. a) after supplied oxygen is consumed, oxygen concentrations immediately start to increase again (0-8h). The oxygen concentration increases over time when no oxygenated water is added (8h-20h). b) cyanide additions lead to a strong increase in oxygen production. Two parallel incubations showed the same pattern as observed in 1a: oxygen increased immediately after added oxygen had been consumed and accumulated over time when no oxygen additions were performed. After the additions of cyanide (0.5mM final concentration, shaded areas), oxygen accumulations strongly increased. Stars: additions of oxygen saturated water, arrows: addition of cyanide. Colors: black and grey lines represent parallel incubations.

In comparison, no oxygen build-up was detected in filtered abiotic controls or when cells were killed by the addition of mercuric chloride (Fig. S1) ruling out the possibility of abiotic oxygen production or intrusion of oxygen into the incubation bottle. Contamination by oxygen intrusion was further ruled out with incubations in an anaerobic chamber showing the same trend of oxygen production (Fig. S2). Involvement of medium components (e.g. HEPES, EDTA) in oxygen production was also excluded (Fig. S3), and furthermore, oxygen microelectrodes, which make use of a different oxygen measurement principle, showed the same patterns of oxygen increase as the optode measurements (Fig. S4). Incubations with *N. maritimus* in medium containing pyruvate showed no difference compared to incubations without pyruvate (Fig. S5). This experiment rules out oxygen production by H_2_O_2_ dismutation (H_2_O_2_ → H_2_ + O_2_) as pyruvate reacts with and removes H_2_O_2_ via an abiotic decarboxylation reaction (*11*).

As described in more detail below, we measured nitric oxide (NO) concentrations with a microelectrode and found that NO slightly interferes with the optode measurements of O_2_ (Fig. S6) but not the O_2_ microelectrode measurements. Therefore, when optode O_2_ concentrations and NO were simultaneously measured, a NO correction was applied to the O_2_ measurements. The correction, however, was relatively small (0-17%) and fully predictable from the NO concentrations. This correction was only applied when NO and O_2_ were simultaneously measured, recognizing that other optode O_2_ measurements may be slight overestimates (depending on the NO concentration) of the actual O_2_ concentration (Fig. S7).

Despite this small interference on our oxygen measurements, *N. maritimus* clearly produces oxygen when the culture reaches anoxia. We hypothesize that oxygen accumulates as a net balance of simultaneous oxygen production and oxygen consumption by ammonia oxidation, where plateauing oxygen concentrations over time represent a balance between these processes. To test this hypothesis, cyanide (0.5 mM) was added to the oxygen-producing culture. Cyanide inhibits oxygen respiration by the heme–copper oxygen reductase, and thus inhibits ammonia oxidation (*12, supplementary information*). Upon cyanide addition, and after an initial lag phase, oxygen concentrations steadily increased at rates ca. 5 times higher (65±12 nmol/L/h) than before cyanide addition (14±2 nmol/L/h) (Fig. 1B). In similar experiments where NO was also measured, O_2_ increase was uncoupled from NO concentration after cyanide addition (Fig. S8). Overall, these results are consistent with the hypothesis that, in the absence of cyanide, some portion of oxygen production by *N. maritimus* is utilized within the cells and does not accumulate into the surroundings.

We tracked the conversion of ^15^N-labelled ammonium to nitrite to directly explore if *N. maritimus* continues to oxidize ammonia while producing oxygen. In this experiment, cell cultures were washed to reduce the high nitrite background that accumulated (ca. 1 mM) during normal aerobic growth. After this, ^15^N-ammonium (I: 5 μM and II: 25 μM) was added as well as a small amount of ^14^N-nitrite to “capture” any produced ^15^N-nitrite from further transformations. The labelling experiments showed continued ammonia oxidation to nitrite during oxygen production (Fig. 2). These results, consistent with the cyanide addition experiments, confirmed that ammonia oxidation occurs together with oxygen production. Furthermore, the rates of ammonia oxidation in these duplicate experiments were 46 (I) and 39 nM/h (II) (table 1), requiring an oxygen production rate of 69 and 60 nM/h respectively, given the stoichiometry of ammonia oxidation (NH_3_ + 1.5O_2_ → NO_2_^-^ + H_2_O + 1H^+^). Oxygen accumulated at an average rate of only 1.2 nM/h in both experiments. Our results further demonstrate that most of the oxygen produced by *N. maritimus* was immediately consumed through ammonia oxidation. The cell density in these incubations was 1.3*10^7^ cells mL-^1^, and therefore, the average ammonia oxidation rate per cell was 3-3.5 amol/cell/h. In marine OMZs with typical AOA cell densities of about 1-10*10^4^ cells mL^−1^ this would correspond to ammonia oxidation rates of 1-10 nM/d, which are in the range of anammox rates in open-ocean oxygen-minimum zones, the so far only known anaerobic ammonia oxidation process that is considered to be relevant in marine OMZs (*8*).

**Fig. 2.**
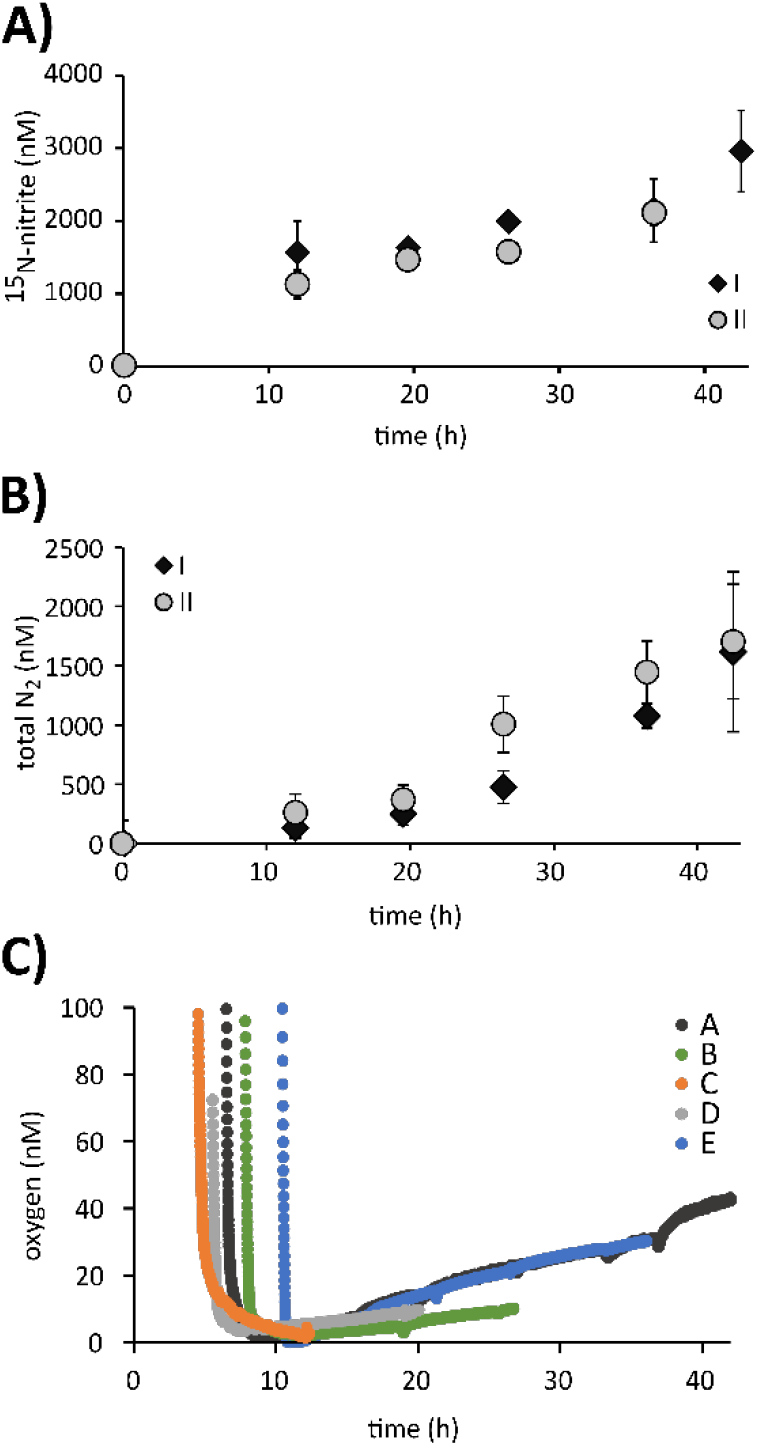
Ammonia oxidation to nitrite and N_2_ during oxygen production by *N. maritimus*. A) ^15^N-nitrite production from ^15^N-ammonium for two sets of incubations of washed *N. maritimus* culture. Incubation I contained a ^14^N-nitrite pool of 5 μM and incubation II had a ^14^N-nitrite pool of 25 μM. ^15^N-nitrite production continued after supplied oxygen was consumed (10h). B) Total N_2_ production in incubations I and II. Results include ^28^N_2_ from ^14^N-nitrite as well as ^30^N_2_ and ^29^N_2_ from added ^15^N-ammonium that was converted to ^15^N-nitrite and partly captured in the small ^14^N-nitrite pool before further conversion to ^30^N_2_ and ^29^N_2_ (results in Fig. S11). C) Oxygen accumulation in a subset of exetainers. Extainers A and B belong to incubation I, and C, D and E to incubation II. Error bars represent the standard deviation of 3 replicates.

**Table 1:**
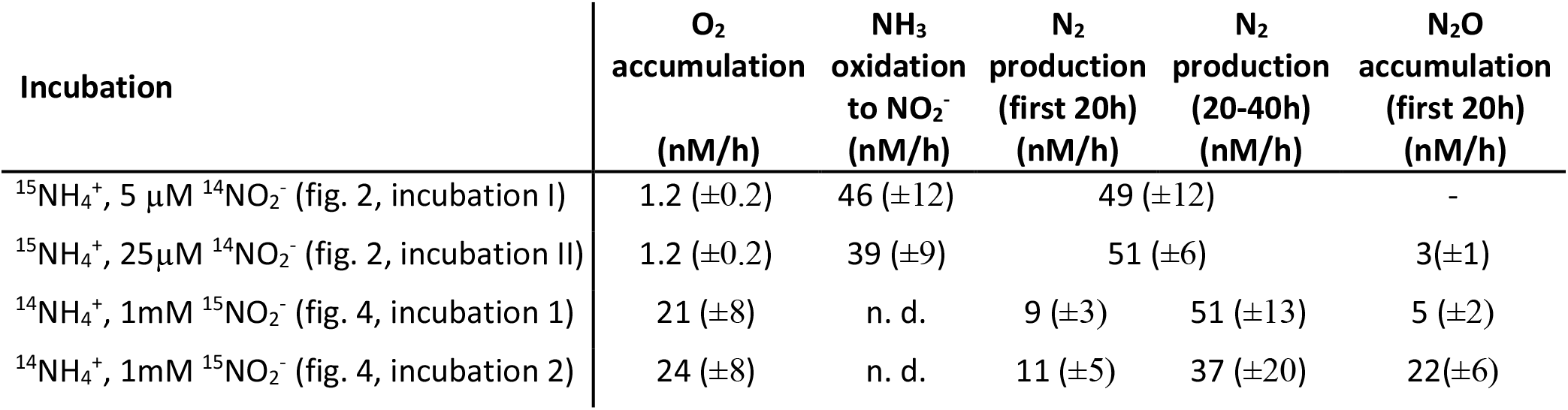
Summary of rates extracted from incubations with ^15^NH_4_^+^ or ^15^NO_2_^-^ additions presented in Figs. 2 and 3. Oxygen accumulation rates were taken when the accumulation rate was at its maximum at the start of oxygen production. In incubations I and II oxygen supplied to the culture at the beginning of the incubation was consumed after 10h (Fig. 2C). Therefore, only time points after 10h were used to calculate ammonia oxidation rates during oxygen production. The means of the rates from replicate incubations and its standard deviation are presented. n. d.: not determined, -: no N_2_O accumulation was detected. N_2_ production rates refer to the total N_2_ production, which in case of incubations 1 and 2 equal ^30^N_2_ production rates.

To explore for translational changes associated with oxygen production, we performed differential proteomic analysis comparing *N. maritimus* performing standard aerobic metabolism to its metabolism during oxygen production. The proteome translation profile during aerobic metabolism matched earlier findings (*14*, see supplementary information for details). However, compared to the mid-log phase during normal aerobic growth, 88 of the 1453 recovered proteins had significantly increased abundances when the culture produced oxygen, while 11 proteins were significantly decreased in abundance (P<0.05; table S1).

The plastocyanin Nmar_1665, the multicopper oxidase type 3 Nmar_1354 and the putative nitroreductase Nmar_1357, were among the most up-regulated proteins during oxygen production, suggesting that they could play a role in the oxygen-production pathway (Fig. S9). Indeed, nitric oxide processing has been suggested for the gene cluster Nmar_1352-1357 (*13, 14*), which would be consistent with the tight coupling of oxygen production and nitric oxide accumulation as explored below. The 17 small blue Cu-containing plastocyanins encoded by *N. maritimus* have a proposed function in hydroxylamine (NH_2_OH) oxidation (*14*). Of these, Nmar_1665 was the only plastocyanin with a significant upregulation under oxygen production, while all other plastocyanins were abundant under both metabolic modes. Other proteins that are either significantly up or down regulated include different transcriptional regulators: e.g members of the AsnC family (Nmar_1524 [up], Nmar_1292 [up], Nmar_1628 [down] (Fig. S9)). Archaeal AsnC regulatory proteins have a function in the regulation of central and energy metabolism, and a role in the switch between aerobic and anaerobic metabolism has been suggested (*15*, *16*). The NADH dehydrogenase 30 kDa subunit Nmar_0278 was significantly down regulated. Otherwise, no proteins with functions in energy or carbon metabolism changed significantly in abundance.

Overall, the few significant changes in the proteome between aerobic respiration and oxygen production suggest that most proteins are needed and active under both metabolic modes. However, some of the upregulated proteins could be involved in catalyzing and regulating oxygen production in AOA.

We now explore possible metabolic pathways for dark oxygen production in *N. maritimus*. Of the three known pathways of dark oxygen production, we rule out perchlorate/chlorate respiration as our cultures were perchlorate/chlorate/chlorite free. Furthermore, as discussed above, we also rule out hydrogen peroxide dismutation as a source of oxygen. As *N. maritimus* metabolizes nitrogen and accumulates NO under normal aerobic ammonia oxidation (*17*), NO dismutation becomes a potential source of oxygen in our experiments. So far, NO dismutation is only known among the NC10 bacteria (*10*). These organisms are methane oxidizers and generate NO for dismutation to oxygen and dinitrogen (*18*), where the oxygen is used to oxidize methane. Because oxygen production and consumption are tightly coupled, methane-oxidizing NC10 bacteria are not known to liberate free oxygen into the environment (*18*).

Using NO microelectrodes, we noted that NO and oxygen production were mostly tightly coupled (Fig. S7) (although the coupling was less tight in other cases; Fig. S8)). Furthermore, when the NO scavenger PTIO (2-Phenyl-4,4,5,5-tetramethylimidazoline-1-oxyl 3-oxide) was added, oxygen production ceased (Fig. S10). Taken together, these results suggest that NO is a crucial intermediate in oxygen production.

We used ^15^N-labelled nitrite to further unravel the pathways of nitrogen and oxygen cycling during oxygen production by *N. maritimus*. In incubations with added ^15^N-nitrite, ^30^N_2_ was produced during oxygen production (Fig 3A, S12 and 13), while no formation of ^29^N_2_ was detected (Fig. 3B). Dinitrogen production by *N. maritimus* or other AOA isolates has not previously been reported. Our results furthermore show that both nitrogen atoms in the N_2_ originated from nitrite, with none coming from ammonium. This result was confirmed by incubations with ^15^N-ammonium, where no immediate conversion of ^15^N-ammonium to ^29^N_2_ or ^30^N_2_ was detected (Fig. 3A, B). Instead, ^15^N-ammonium was most likely converted to nitrite and diluted into the large existing nitrite pool in this experiment. In contrast, when ^15^N-labelled ammonium was added to washed cultures with a small nitrite pool (5 and 25 μM), the ^15^N-nitrite produced from ammonia oxidation was further converted to N_2_ (Fig. 2B and S11) consistent with our experiments with ^15^N-labelled nitrite.

**Fig. 3.**
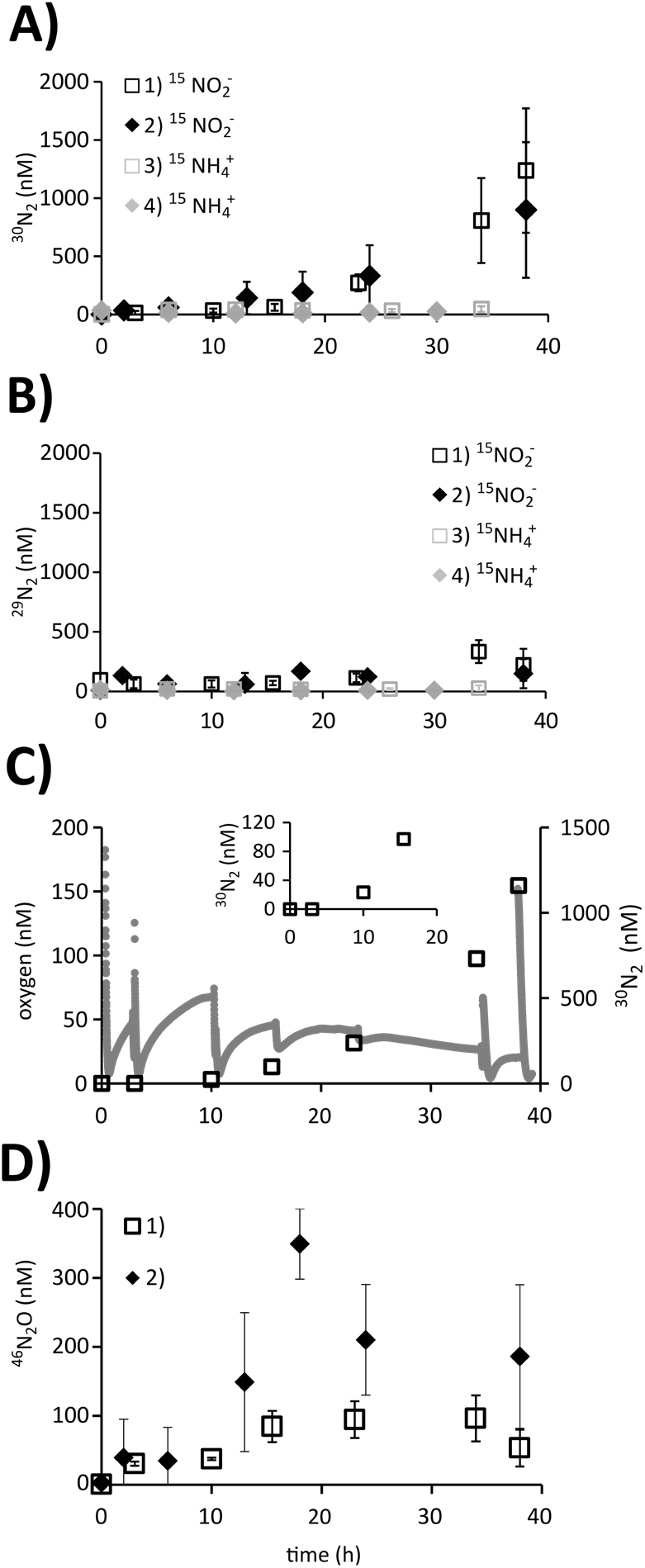
N_2_ production and N_2_O accumulation by *N. maritimus* during simultaneous oxygen production. After sparging with argon, *N. maritimus culture* was incubated with either ^15^N-nitrite or ^15^N-ammonium and the production of (A) ^30^N_2_ and (B) ^29^N_2_ were tracked in two independent sets of incubations for each tracer addition (3-4 replicates each). N_2_ production was only detected in incubations with ^15^N-nitrite. For the corresponding oxygen measurements see fig. S12 and 13. C) Oxygen accumulation and ^30^N_2_ production for a single replicate of the incubation series 1). The insert shows ^30^N_2_ production in the first 20h. Disturbances in the oxygen time series at T=0, 3, 10, 15.5, 23, 34 and 38h correspond to the time points when samples for N_2_ analysis were taken, which led to slight oxygen intrusion. Gray dots: oxygen, black open squares: ^30^N_2_. D) ^46^N_2_O accumulation in incubations 1) and 2) supplied with ^14^N-ammonium and 1 mM ^15^N-nitrite. Only ^46^N_2_O accumulated in these incubations with a large 15N-nitrite pool indicating that all produced N_2_O originated from nitrite. Error bars represent the standard deviation of 4 (incubations 1 and 3) or 3 (incubations 2 and 4) replicates.

Thus far we have shown that NO is a likely intermediate in oxygen production and that N_2_ is produced together with O_2_ by *N. maritimus*. Rates of O_2_ accumulation and N_2_ production from the different incubations shown in Figs. 2 and 3 are summarized in table 1. These results would generally be consistent with NO dismutation as a source of both N_2_ and O_2_, where NO is produced from the reduction of nitrite. In incubations with added ^15^N-nitrite, however, oxygen accumulation exceeded N_2_ production in the first 20 h (incubations 3 and 4 in Table 1, Fig. 3c, S12 and S13), demonstrating a decoupling of O_2_ and N_2_ production in this phase of the experiment. As net rates of O_2_ accumulation may underestimate gross rates of O_2_ production, as explored above, and as N_2_ production in our experiments is a gross production rate, there is an imbalance between the production rates of O_2_ and N_2_. Such an imbalance would be inconsistent with O_2_ and N_2_ production directly from NO dismutation. In this case the O_2_ accumulation rates should not exceed N_2_ production rates, but the imbalance we measure suggests that further intermediate(s) must exist between NO, and N_2_ and O_2_ production. We suggest that N_2_O may be such an intermediate, where 2NO → N_2_O + 0.5O_2_.

Indeed, in our incubations supplied with ^15^N-nitrite, ^46^N_2_O accumulated before ^30^N_2_ production accelerated (Fig. 3D), and the rates of N_2_ production and N_2_O accumulation taken together in the first 20 h match the O_2_ accumulation rates within the uncertainties (table 1). Furthermore, the dismutation of NO(aq) to O_2_(aq) and N_2_O(aq) is thermodynamically favorable (ΔG^0^’= −165 KJ/mol O_2_,). However, for this pathway to occur, an unknown N_2_O reductase would need to be present in the genome of *N. maritimus* (*14*). N-nitrosating hybrid formation, in which one N atom from NO_2_^-^ and one from NH_4_^+^ (or an intermediate of ammonia oxidation) combine to form N_2_O, has been proposed as a possible source for N_2_O production in AOA (*19*), but the isotopic signature of the N_2_O produced in our experiments (^46^N_2_O) does not support this pathway (expected ^45^N_2_O). In incubations supplied with ^15^N-ammonium and a small nitrite pool (Fig. 2), N_2_O accumulated transiently as well (Fig. S14). In these incubations N_2_ production far exceeded O_2_ accumulation and no N_2_O accumulation would be required for a mass balance. This is not surprising as high rates of ammonia oxidation (Table 1, Fig 2) indicate that O_2_ accumulation rates in these experiments far underestimate gross rates of O_2_ production as explored above. This does not mean that N_2_O was not an intermediate in these experiments, only that these incubations did not demonstrate an initial imbalance between O_2_ and N_2_ production.

To summarize, like for NO dismutation in NC10 bacteria, O_2_ production in *N. maritimus* has NO as an intermediate and produces N_2_. However, unlike for NC10 bacteria, our results suggest that the pathway of O_2_ production employed by *N. maritimus* has an extra intermediate that may be N_2_O. A proposal for the metabolic pathway associated with oxygen production in *N. maritimus* is shown in figure 4. While ammonia oxidation to nitrite is accomplished by the O_2_ produced by *N. maritimus*, the conversion of nitrite to N_2_ requires reducing equivalents regardless of the O_2_ production pathway. The required electrons can partly, but not fully, be obtained from the ongoing oxidation of ammonia. Another source of electrons could be intra- or extracellular organic matter produced during normal aerobic ammonia oxidation (*20*) or, *in situ*, by dissolved organics available in the water column.

**Fig. 4.**
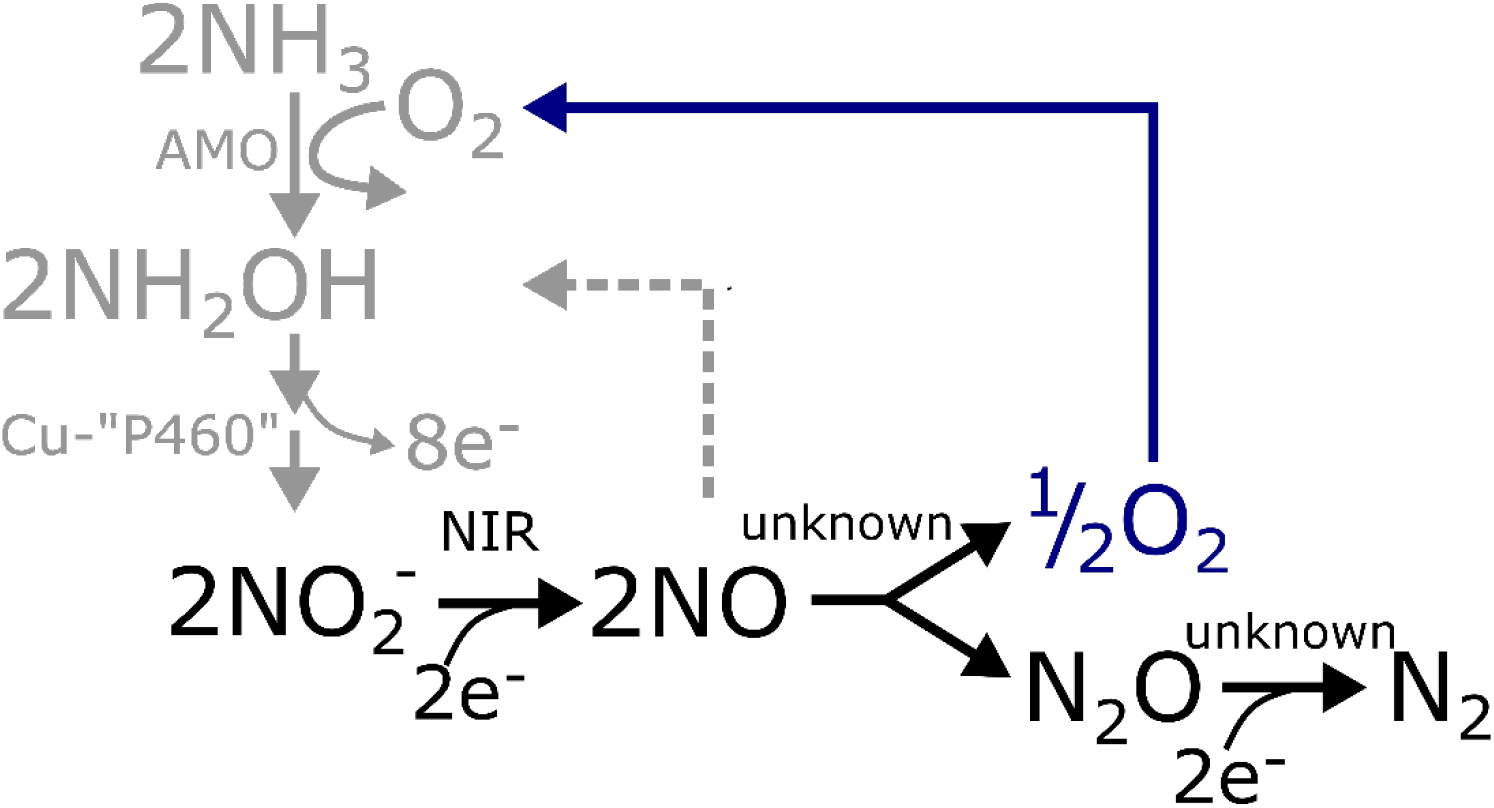
Proposed pathway of oxygen and dinitrogen production in *N. maritimus*. Ammonia oxidation pathway (gray): for the aerobic oxidation of ammonia to nitrite, oxygen is needed to activate ammonia oxidation by the ammonia monooxygenase (AMO), additionally four electrons per oxidized NH_4_^+^ are transferred to the terminal oxygen reductase, which reduces oxygen. Proposed oxygen production pathway (black): Nitrite is reduced to nitric oxide by the NirK nitrite reductase (NIR). Nitric oxide is dismutated to oxygen and nitrous oxide. The accumulating oxygen is partly consumed during ammonia oxidation. Likewise, part of the nitric oxide may be used for driving the hydroxylamine oxidation step of the ammonia oxidation pathway. Finally, nitrous oxide becomes reduced to dinitrogen. This pathway requires four electrons per produced N_2_. These may partly be supplied by the ammonia oxidation reaction, which in return would reduce the oxygen demand by the ammonia oxidation pathway. Cu-“P460”: the hydroxylamine oxidizing enzyme, unknown: unknown enzyme.

By showing that the O_2_ production pathway in AOA is coupled to N_2_ production, we also uncovered a new, potentially environmentally significant, pathway of N_2_ production. Furthermore, by converting ammonium through nitrite to N_2_, AOA perform a so far unrecognized pathway of anaerobic ammonia oxidation. ^15^N-tracer experiments currently performed to measure N-cycling rates in marine oxygen-depleted environments would overlook this pathway and account for its N_2_ production as canonical denitrification and/or anammox.

AOA are one of the most abundant groups of microbes in the global ocean and key players in the marine nitrogen cycle, also in low-oxygen environments. Thus, a widely distributed oxygen-producing pathway by AOA could have far-reaching implications for the microbial ecology and biogeochemical cycling in oxygen-depleted environments, including the possibility that some of the produced could be used by other microbial cells. The discovery of an anaerobic oxygen-producing pathway in AOA can explain the presence and role of AOA in such environments solving a longstanding enigma.

## Acknowledgments

We thank Annie Glud for assistance with microelectrode measurements and providing microelectrodes.

## Autor contributions

B.K. and D.E.C. designed the experiments. B.K. performed the experiments and analyzed data with input from M.L, L.B., M.K., B.T. and D.E.C.. N.J. performed proteomics. B.K. and D.E.C. wrote the manuscript, with contributions and approval from all other authors.

## Data and materials availability

All data is available in the main text or the supplementary materials.

## Supplementary Materials

Materials and Methods

Figures S1-S14

Tables S1-S2

References (21-30)

