## supplementary information for "Oxygen production by an ammonia-oxidizing archaeon"

**This PDF file includes:**

Materials and Methods
Supplementary Text
Figs. S1 to S14
Captions for Data tables S1 and S2

**Other Supplementary Materials for this manuscript include the following:**

Data tables S1 and S2

### Material and Methods.

Culturing of *N. maritimus* SCM 1

*N. maritimus* SCM 1 was cultured aerobically at 28°C in HEPES-buffered synthetic Crenarchaeota medium containing (/L): NaCl [26 g], $MgCl_2x6H_2O$ [5 g], $MgSO_4x7H_2O$ [5 g], $CaCl_2$ [1.5 g] and KBr [0.1 g], 10 ml HEPES buffer [1 M HEPES at pH 7.8], 2 ml $NaHCO_3$ [1 M], 5 ml $KH_2PO_4$ [0.4 g/l], 1 ml EDTA complexed trace element solution (*21*) and 1 mM $NH_4Cl$ (*9*, *22*). The pH of the medium was adjusted to 7.5.
To exclude any effect (e.g. interference) of supplied organic carbon components in the medium, controls were performed with *Nitrosopumilus maritimus* SCM 1 cultured in HEPES-free medium (6 ml $NaHCO_3$ [1 M]) and 1ml of an EDTA-free acidic trace element solution was added (*21*). Prior to incubations cultures were checked microscopically for potential microbial contaminants.

Incubations

Incubations were performed in custom-modified Schott Duran glass bottles of 1160 mL as described in Tiano *et al*. 2014 (*7*). These bottles have three openings: A 25 cm long glass capillary (internal diameter 0.25 cm), a microelectrode insertion port and a modified opening at the bottle neck closed with a ground glass stopper during incubations. The bottles were designed so that diffusion of oxygen into the bottle is negligible (*7*). Optode spots for oxygen measurements (see below) were glued to the inside of each bottle. Prior to incubation, the culture bottles were purged with argon for 40 min to lower the oxygen concentration in the culture. The incubation bottles were then filled from the culture bottle through a glass tube connection (with Viton tubing joints) to the incubation bottles and by applying overpressure to the culture bottle with argon gas flow. A solid glass bar with the same dimension as the microelectrodes was inserted into the microelectrode insertion port when a microelectrode was not used (*7*). Bottles were stirred constantly with a glass-coated magnet (45 mm) at 400 rpm. Bottles, glass-coated magnets and glass inserts had been autoclaved before use.

The incubation bottles were kept in the dark in a temperature-controlled water bath at a constant 28°C. In order to measure oxygen respiration kinetics, and to calibrate the oxygen optodes (see below), defined amounts of oxygen-saturated MilliQ water were added at the beginning or the end of the incubation period. Dissolved substrates, oxygenated water, and inhibitors were added by a glass syringe connected to a metal needle reaching through the capillary into the bottle.

$N_2$ production

To determine the pathways of $N_2$ production during $O_2$ production by *N. maritimus* incubations with either $^{15}$N-nitrite (incubations 1 and 2) or $^{15}$N-ammonium (incubations 3 and 4) were performed:

$N_2$ production via nitrite reduction (incubations 1 and 2)

To test for nitrite reduction during oxygen production by *N. maritimus*, freshly inoculated cultures received $^{15}$N-ammonium (1 mM) as substrate that they oxidized to $^{15}$N-nitrite during aerobic growth. After complete conversion to $^{15}$N-nitrite (monitored spectrophotometrically), 500 µM $^{14}$N-ammonium was added, and incubations were performed in glass bottles as described above.

Samples for nutrient analyses and for the analysis of the isotopic compositions of $N_2$ and $N_2O$ were taken with a glass syringe connected to a metal needle reaching through the capillary. At the same time the culture volume was sampled, it was replaced by anoxic medium injected by another glass syringe and metal needle. Samples for $N_2$ and $N_2O$ analysis were transferred to 3 ml gas-tight glass vials (Extainer, Labco) and fixed with 0.5 ml of zinc chloride (50% w/v). Nutrient samples were immediately frozen until further analysis.

$N_2$ production from ammonium (incubations 3 and 4)

To test for the conversion of ammonia to $N_2$ during oxygen production by *N. maritimus,* $^{15}$N-labelled ammonium (100-500 µM) was added to incubations with cultures that had previously accumulated 1mM of $^{14}$N-nitrite during their aerobic growth. The cultures were transferred to glass bottles and incubations were performed as described above. Samples for the analysis of the isotopic composition of $N_2$ and $N_2O$ analysis were transferred to exetainers and treated as described above.

<u>Control incubations</u>

Further control incubations were performed with the following changes from standard procedure:

1. *N. maritimus* was grown in medium without HEPES and EDTA to exclude abiotic oxygen production by interaction of the medium components with intermediates and products of *N. maritimus* physiology. After several transfers of the culture in HEPES and EDTA-free media, culture incubations in the HEPES and EDTA-free media were performed in glass bottles as described above.
2. *N. maritimus* was grown in medium containing 0.2 mM pyruvate as described above in order to exclude $H_2O_2$ dismutation as a potential oxygen production mechanism as α-Keto acids detoxify $H_2O_2$ via an abiotic decarboxylation reaction (*11*).
3. Potassium cyanide (0.5 mM) was added to incubations in order to inhibit oxygen respiration by the heme–copper oxygen reductase and to quantify net oxygen production.
4. The NO-scavenger PTIO was added to mid-incubation to a final concentration of 100µM in order to test if NO is an intermediate in oxygen production.
5. For killed controls, saturated mercuric chloride solution was added to incubations of *N. maritimus*.

<u>Ammonia oxidation (incubations I and II)</u>

During aerobic growth, *N. maritimus* cultures accumulated high concentrations of nitrite through ammonia oxidation. To measure rates of ammonia oxidation to nitrite in incubations with $^{15}$N-labelled ammonium during oxygen production, this nitrite background needed to be reduced. Therefore, *N. maritimus* cultures were washed with ammonium-free medium and the cells concentrated using Vivaspin 20 centrifugal concentrator columns (100kDa; Sartorius, Germany). Washed cells were pooled from several columns. Nitrite (5µM or 25µM) and $^{15}$N-ammonium (50 µM) was added. Due to the small volume of the washed cell culture, incubations for ammonia oxidation were performed in 3 ml gas-tight glass vials (Extainer, Labco, UK). Optode spots were glued to the bottom of a subset of exetainers while exetainer lids had been kept under helium atmosphere for 3 months prior to the incubations to reduce a source oxygen contamination from the lids (*23*).

Washed cells were sparged with helium for 30 min and then transferred to the exetainers with a glass syringe. To avoid diffusion of small amounts of oxygen through the exetainer lids during the incubation time, incubations were performed in an anaerobic chamber (Coy Laboratory Products, ~2% $H_2$). To stop the activity in the exetainers at the respective time points, the cells were killed directly in the exetainers by adding 0.5 ml of zinc chloride (50% w/v); 0.5 ml of sample was discarded simultaneously. The culture liquid in the preserved exetainers was used for the determination of nitrite concentrations and the isotopic composition of nitrite and $N_2$.

Chemical analyses

Oxygen concentrations (measuring range: 1 - 1000 nM) were measured using a Luminescence Measuring Oxygen Sensor (LUMOS) reading oxygen quenching on optode spots (*6*). The optode spots were formed as glass dots coated with a thin layer of the fluorophore (palladium(II) 5,10,15,20-tetrakis- (2,3,4,5,6- pentafluorphenyl)- porphyrin)) in Hyflon AD 60, and these were glued to the inside of the glass incubation vessels. The LUMOS readout device was mounted to the glass vessel by a PVC holder which was glued to the outside of the bottle aligning with the optode spot (24). The LUMOS readout device was connected to a computer. Data were logged by the software FireSting Logger (Pyroscience).

We determined the interference of NO with optode oxygen measurements by measuring the response of the optodes when different quantities of nitrite were added to anoxic acidic water (purged with $N_2$, pH=1.5), where the nitrite formed NO once dissolved. The concentrations of NO were measured with a NO-500 NO-microelectrode (Unisense, Aarhus, Denmark). In control experiments complementary to LUMOS oxygen detection, a potentiometric oxygen microelectrode was inserted into the incubation bottle (*25*). No interference of NO with the potentiometric oxygen microelectrode was found when tested as described above. The microelectrode currents for both NO and potentiometric $O_2$ determination were measured by a picoammeter PA 2000 (Unisense, Aarhus, Denmark).

Nitrite and ammonium were measured spectrophotometrically according to Bendschneider and Robinson (1952) and Bower and Holm-Hansen (1980) (*26*, *27*). Nitrogen isotopes in nitrite, $N_2$ and $N_2O$ were analyzed by coupled gas chromatography–isotope ratio mass spectrometry (GC-IRMS) on a Thermo Delta V Plus isotope ratio mass spectrometer as described by Dalsgaard *et al.* (2012) (*28*).

Proteomics

For proteomics, *N. maritimus* SCM 1 was cultured aerobically and incubations bottles were set-up as described above. Subsequently, *N. maritimus* cultures were incubated without any oxygen addition for more that 36h, and oxygen production was monitored as described above. One hundred mL of the both the aerobic culture and the oxygen-producing incubations were filtered onto separate PTFE filters (Advantec) and flash frozen in liquid nitrogen. Filtration of the oxygen-producing cultures was performed in the anaerobic chamber. Filters were frozen until further processing.

Filters were cut in small pieces and resuspended in 1 ml lysis-buffer (0.29% NaCl, 0.01M Tris-HCl, 5mM EDTA, 0.4% SDS) with 1 µl PMSF solution. Cells were disrupted by bead beating (FastPrep-24, MP Biomedicals, Sanra Ana, CA, USA; 5.5 ms, 1 min, 3 cycles) followed by ultra-sonication (UP50H, Hielscher, Teltow, Germany; cycle 0.5, amplitude 60%) and centrifugation (10,000 x g, 10 min). The protein lysate was load on SDS-gel and run for 10 min. The gel piece was cut, washed and incubated with 25 mM 1,4-dithiothreitol (in 20 mM ammonium bicarbonate) for 1 h and 100 mM iodoacetamide (in 20 mM ammonium bicarbonate) for 30 min detained, dehydrated and proteolytically cleaved overnight at 37 °C with trypsin (Promega). The digested peptides were extracted and desalted using ZipTip-µC18 tips (Merck Millipore, Darmstadt, Germany).

The peptide lysates were then re-suspended in 0.1% formic acid and injected to nanoliquid chromatography mass spectrometry (nanoLC-MS/MS). Mass spectrometric analysis of peptides was performed on a Q Exactive HF mass spectrometer (Thermo Fisher Scientific, Waltham, MA, USA) coupled with a TriVersa NanoMate (Advion, Ltd., Harlow, UK) source in LC chip coupling mode. LC gradient, ionization mode and mass spectrometry mode were used as described in (*29).*

Data from the LC-MS/MS were analyzed using the Proteome Discoverer program (v.1.4, Thermo Fischer Scientific, Waltham, MA, USA) using SEQUEST HT. As the reference database, the protein-coding sequences of *N. maritimus* were used. Search settings were set to trypsin (Full), max. missed cleavage: 2, precursor mass tolerance: 10 ppm, fragment mass tolerance: 0.02 Da. The false discovery rates (FDR) were determined with the node Percolator (*30*) embedded in Proteome Discoverer, and we set to the FDR threshold at a peptide level of <1%. The same threshold was set for the protein FDR (<1%).

### Supplementary text

Cyanide inhibition

During ammonia oxidation oxygen is the terminal electron acceptor, and in *N. maritimus*, oxygen is reduced by a cytochrome aa3 oxidase. Oxygen is also needed for the activation of ammonia monooxygenase. Cyanide inhibits cytochrome aa3 oxidases by binding to the binuclear cytochrome a3/Cu center of the partially reduced enzyme species formed during turnover *(12)*. Similarly, cyanide is expected to at least partially inhibit the binuclear copper site of the ammonia monooxygenase. However, ammonia monooxygenase inhibition by cyanide has so far only been shown for the bacterial ammonia monooxygenase *(31,32)*. Therefore, inhibitory concentrations are unknown for *N. maritimus*. But cyanide should shut down the biggest sink of oxygen, the terminal oxidases. We cannot exclude that cyanide interferes with other enzymes involved in ammonia oxidation or oxygen production, which may explain the lag phase in oxygen production after cyanide addition.

### Differential proteomics

The most abundant proteins under both normal aerobic growth and oxygen-producing conditions included the predicted surface-layer (S-layer) proteins Nmar_1547 and Nmar_1201, the two putative NirK nitrite reductases Nmar_1667 and Nmar_1259, the ferredoxins Nmar_0239 and Nmar_1537 and the putative oxidoreductase Nmar_1622 and Nmar_1109 (table S2). All proteins deemed important for ammonia oxidation were abundant with no significant differences between metabolic modes. These included the ammonia monooxygenase subunit B (Nmar_1503), complex III (Nmar_1542-1544), complex IV (Nmar_0182-0185), the two NirK paralogs, cupredoxin Nmar_1307, and ammonium transporter AMT2 (Nmar_1698). Only AmoA and AmoC (both present) were poorly recovered, which can be attributed to the digestion method used here (*13*).

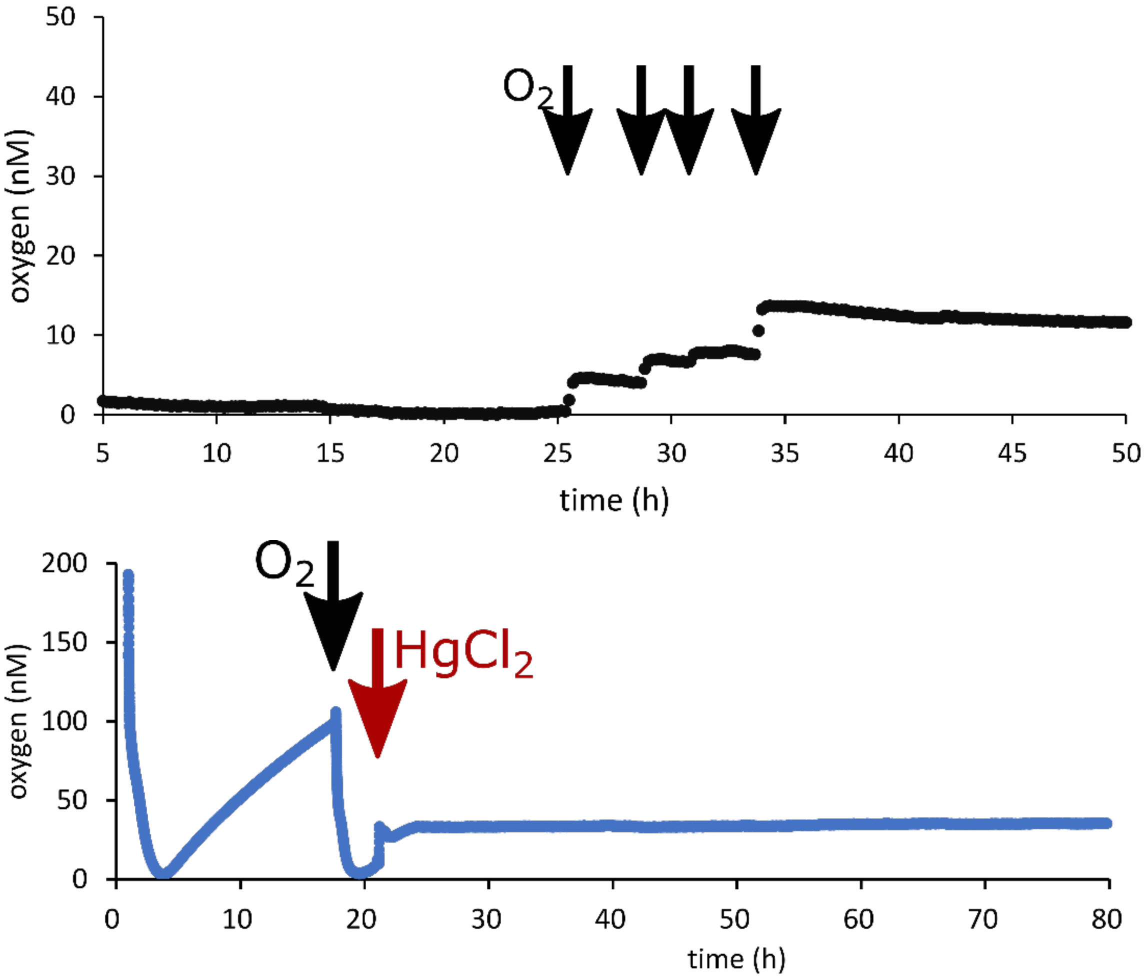


Fig. S1. Controls testing for abiotic oxygen production or intrusion during incubation. A) Abiotic control showing oxygen concentrations hovering near 0 nM for 24h until oxygen was added. After oxygen additions, oxygen concentrations remained stable at the respective new level. B) Normal oxygen production by *N. maritimus* over 21 hours including a small addition of oxygen-saturated water at 18 hours (black arrow). Mercuric chloride was added at 22 hr (red arrow), after which oxygen concentrations remained stable.

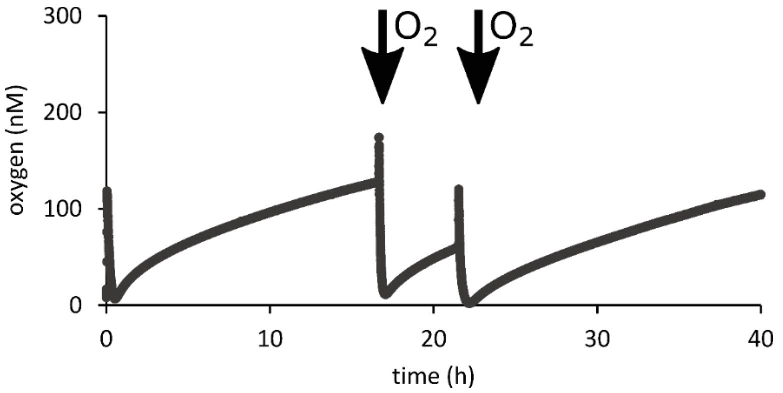


Fig. S2. Oxygen production during incubation of *N. maritimus* culture in an anaerobic chamber. Once the supplied oxygen was completely consumed, oxygen accumulated again at rates similar to those observed in incubations outside the anaerobic chamber (fig 1). Black arrows: additions of oxygen-saturated water.

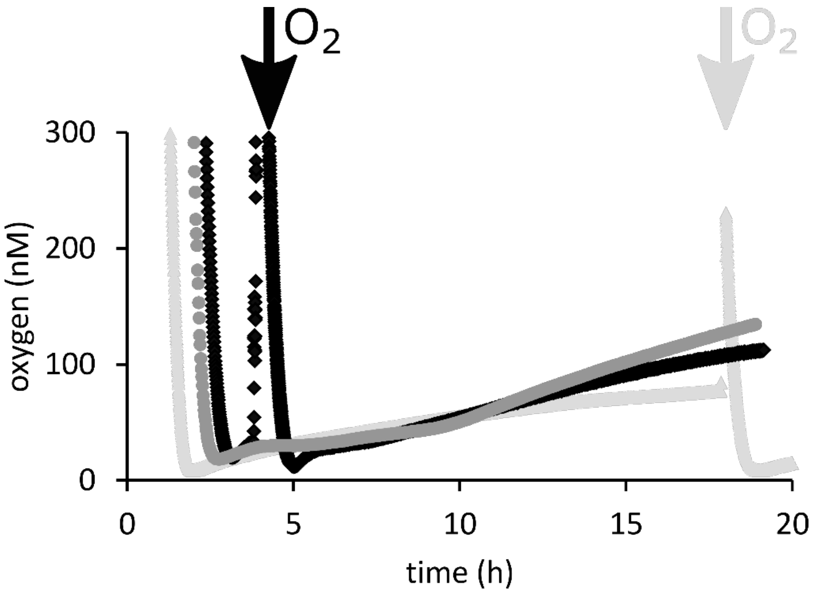


Fig. S3. Time course of the oxygen concentrations in control incubations with *N. maritimus* cultures grown in medium without HEPES or EDTA. Incubations were performed in order to exclude abiotic oxygen production by interaction of the medium components with intermediates and products of *N. maritimus* physiology. Three parallel incubations are depicted in light gray, dark gray and black. Arrows in the corresponding colors indicate the additions oxygen saturated water.

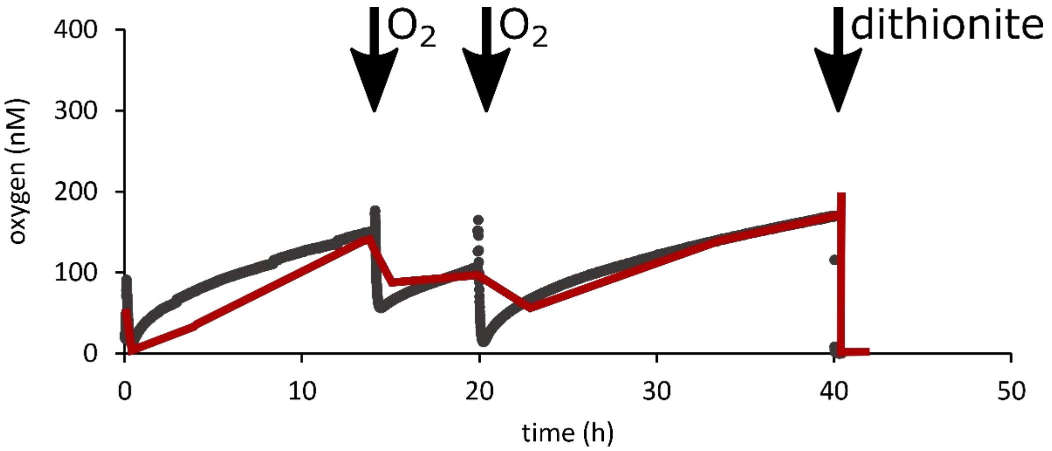


Fig. S4. Oxygen production by *N. maritimus* measured with a LUMOS optode and simultaneously with an oxygen microelectrode. Red: oxygen microelectrode, black: Lumos optode. Black arrows: additions of oxygen-saturated water and dithionite as indicated.

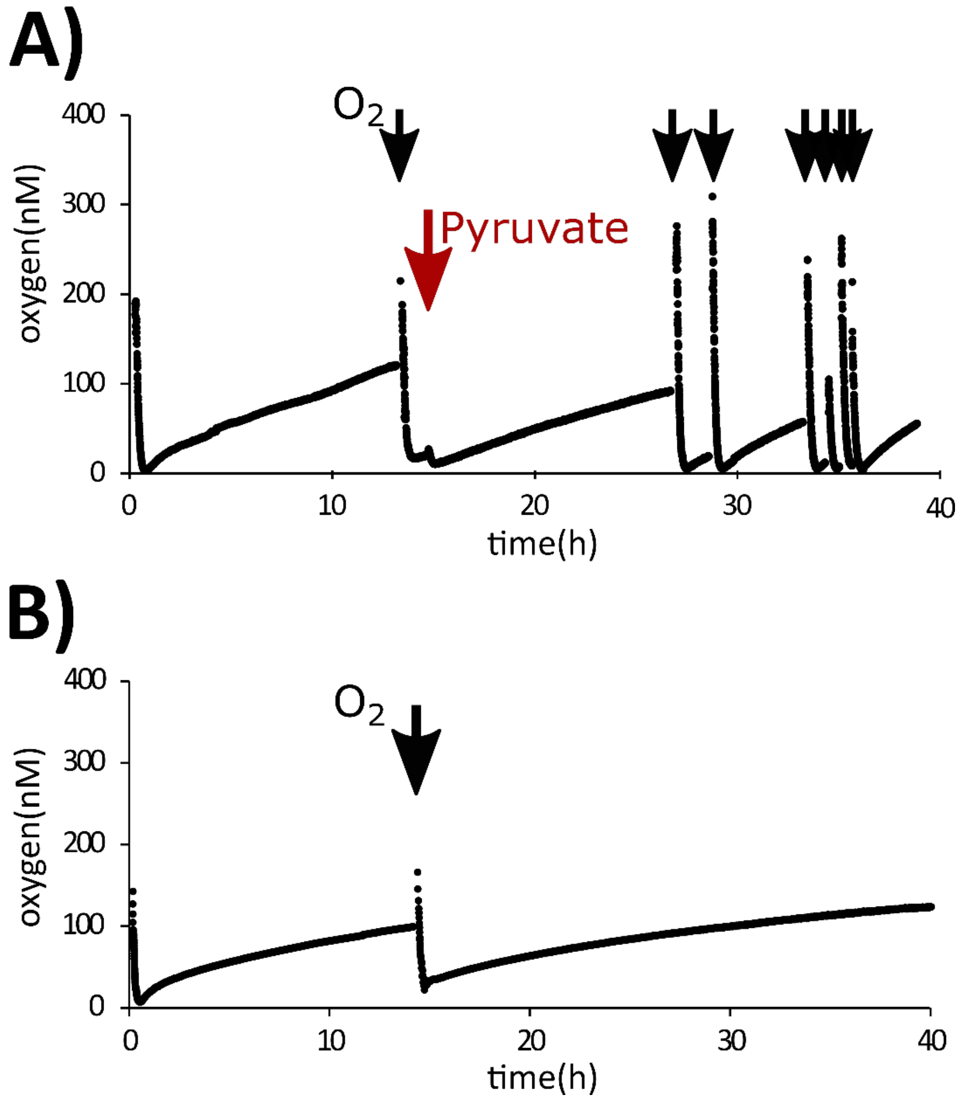


Fig. S5. Time course of oxygen concentrations of incubations with added pyruvate. A) The addition of 0.5 mM pyruvate (final concentration) at hour 14 (red arrow), does not influence oxygen dynamics in oxygen-producing *N. maritimus* incubations, B) Oxygen increases over time in incubations with *N. maritimus* culture grown in a medium containing 0.2 mM pyruvate. Black arrows: additions of oxygen-saturated water.

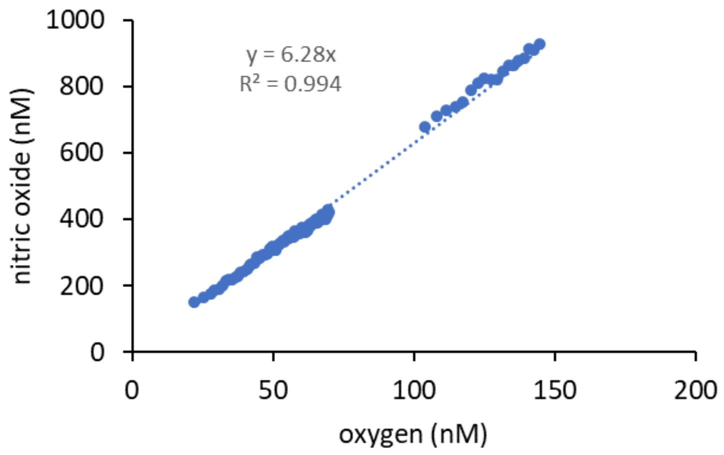


Fig. S6. Interference of NO with oxygen measurements by the LUMOS optodes. Nitrite was converted to NO in an acidic solution, with NO concentrations monitored with a microelectrode and the oxygen response monitored with a LUMUS optode. The oxygen optode response was due to interference by NO. The resulting linear relationship between NO concentration and optode response was used to correct for NO interference when both NO and oxygen concentrations were measured in the same incubation.

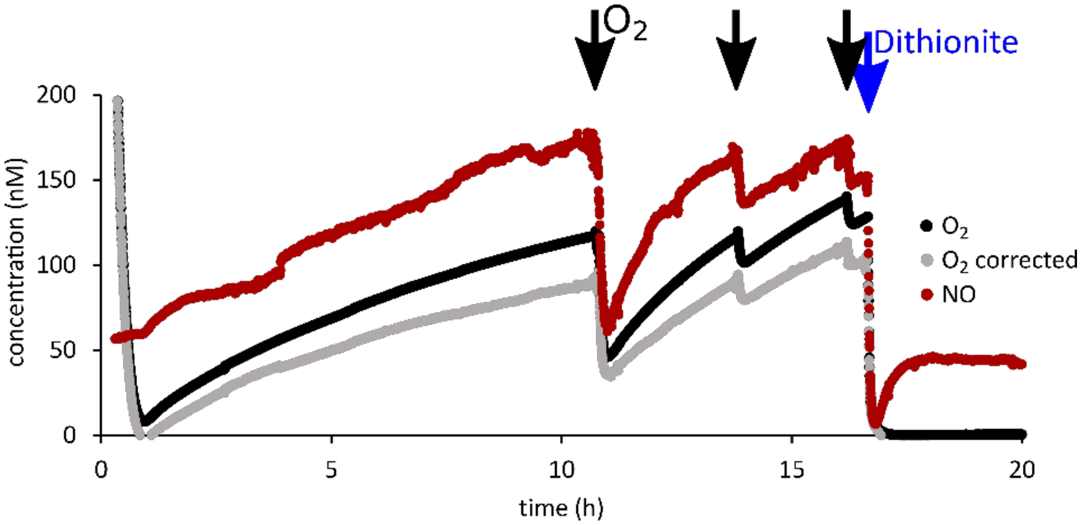


Fig. S7. Incubation of *N. maritimus* with simultaneous measurements of oxygen and NO concentrations. Oxygen concentrations were corrected for the interference of NO with the oxygen optodes (see fig. S6). Oxygen and NO production were closely coupled until dithionite (blue arrow) was added near the end of the incubation.

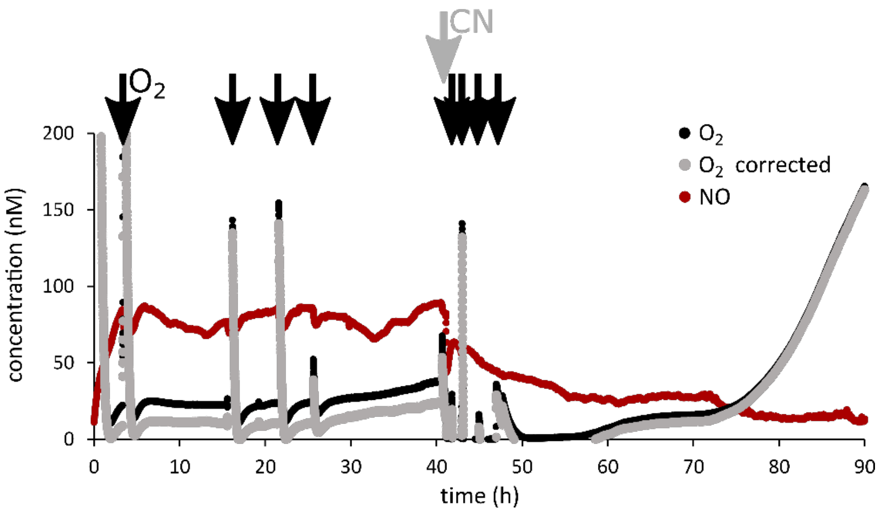


Fig. S8. Incubation of *N. maritimus* with simultaneous measurements of oxygen, oxygen corrected for NO and NO before and after the addition of cyanide (0.5 mM).

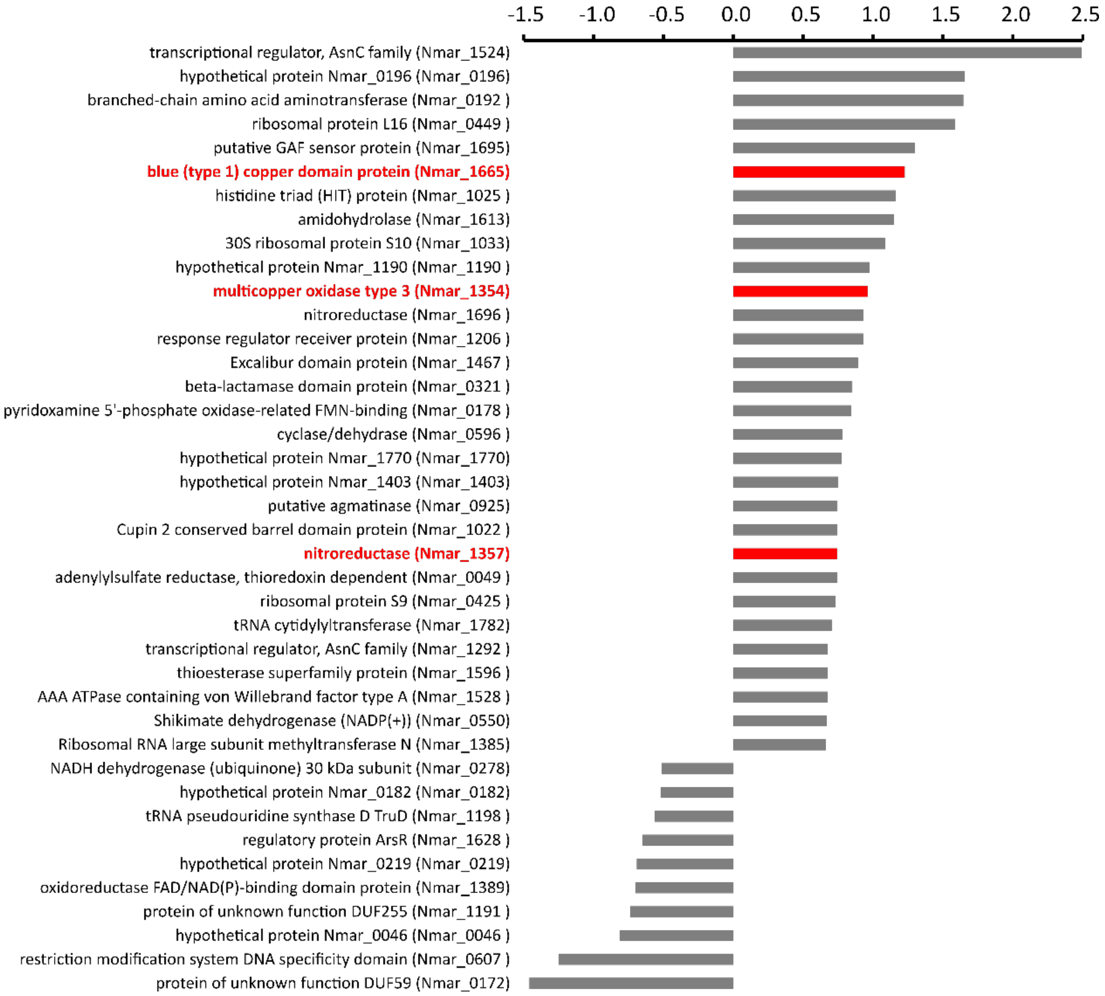


Fig. S9. Changes in protein expression *N. maritimus* during oxygen production compared to when they oxidize ammonium aerobically. $Log_2$ ratio of the 40 proteins with the highest significant increase or decrease in abundance ($P<0.05$). Red: potential involvement in novel oxygen production pathway.

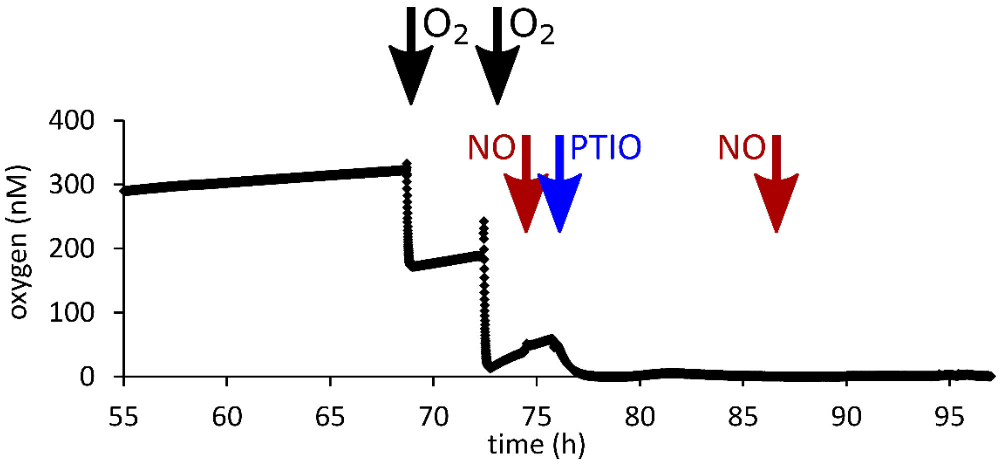


Fig. S10. The production of oxygen in cultures of *N. maritimus* before and after the addition of NO (25 nM final concentration, red arrows) and PTIO (100 µM final concentration, blue arrows). Also shown is the addition of small amounts of oxygen saturated water (black arrows). Before this experiment, oxygen had accumulated to approximately 300 nM over 55h.

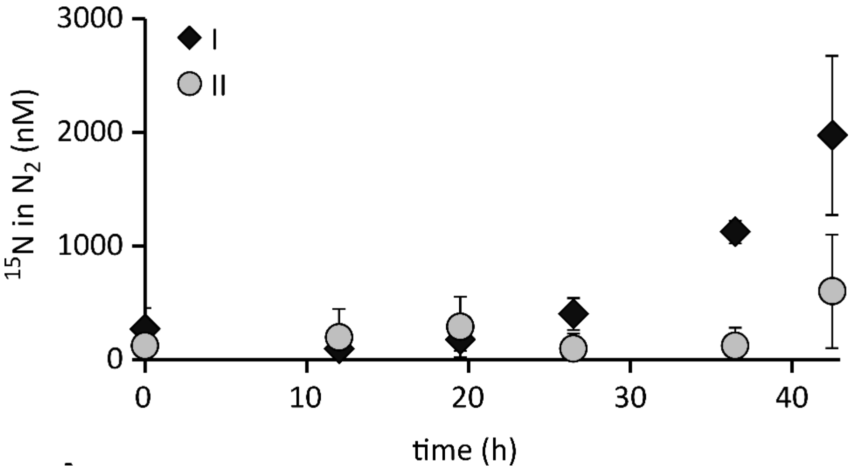


Fig. S11: $^{15}N$ in $N_2$ produced in the incubations I and II (Fig. 2). supplied with $^{15}N$-ammonium and 5 µM (I) or 25 µM (II) $^{14}N$-nitrite. $^{15}N$-ammonium oxidized to $^{15}N$-nitrite is partly captured in the small $^{14}N$-nitrite pool (Fig. 2A) before it is further converted to $^{30}N_2$ and $^{29}N_2$ (paired with $^{14}N$-nitrite). Here, the total $^{15}N$ in the produced $N_2$ is shown in order to demonstrate the conversion of $^{15}N$-ammonium to $N_2$ via nitrite. The total $N_2$ production including $^{28}N_2$ produced from $^{14}N$-nitrite is shown in Fig. 2b.

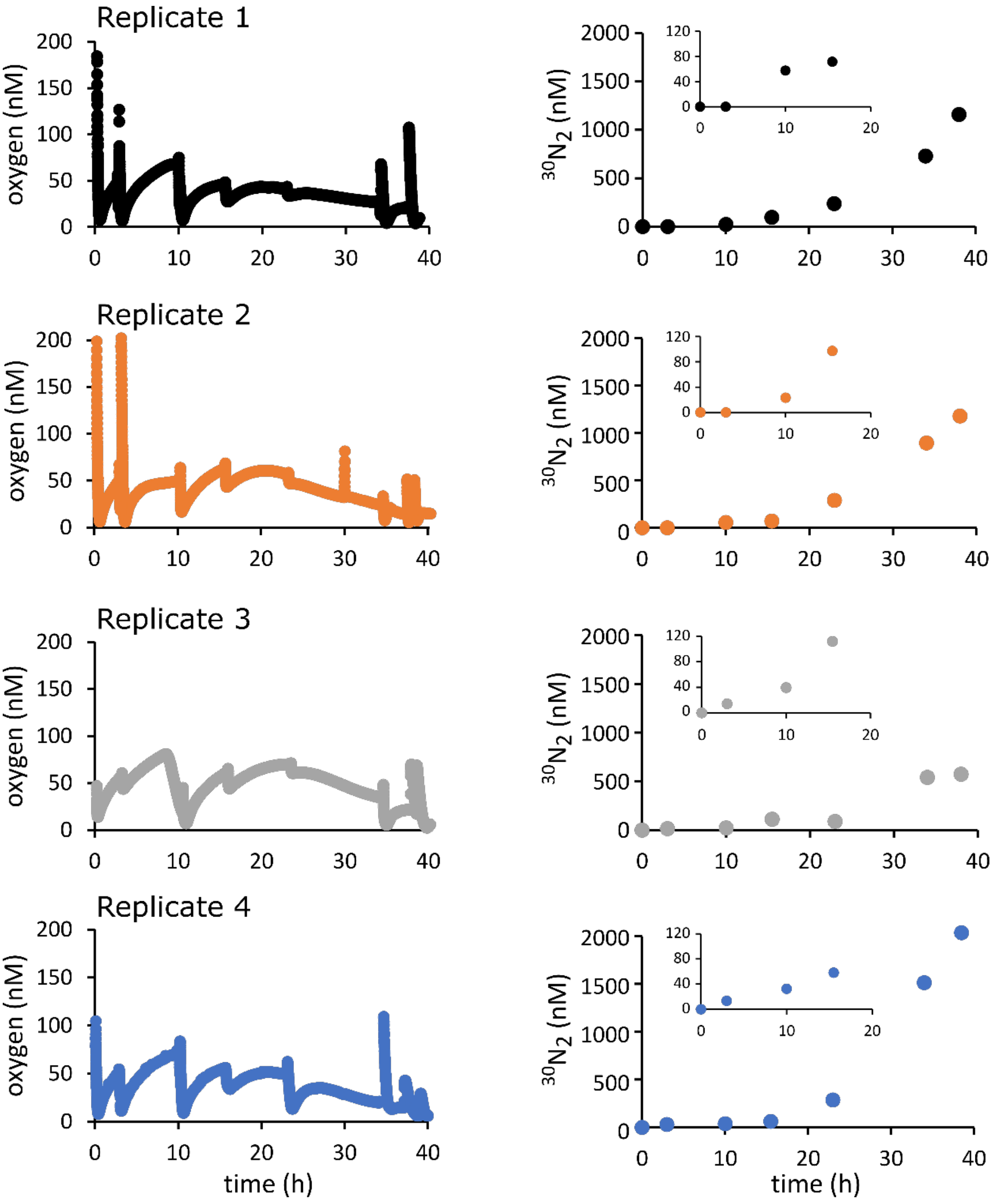


Fig. S12. Oxygen accumulation (left) and $^{30}N_2$ production from $^{15}$N-nitrite (right) in the single replicates of the incubation series 1) in fig. 3 (black open squares). The inserts show the $^{30}N_2$ production in the first 20h of the incubation. Disturbances in the oxygen time series at T=0, 3, 10, 15.5, 23, 34 and 38h correspond to the time points when samples for $N_2$ analysis were taken, which led to slight oxygen intrusion. Only $^{30}N_2$ production is shown as no production of $^{29}N_2$ was detected (see fig 3).

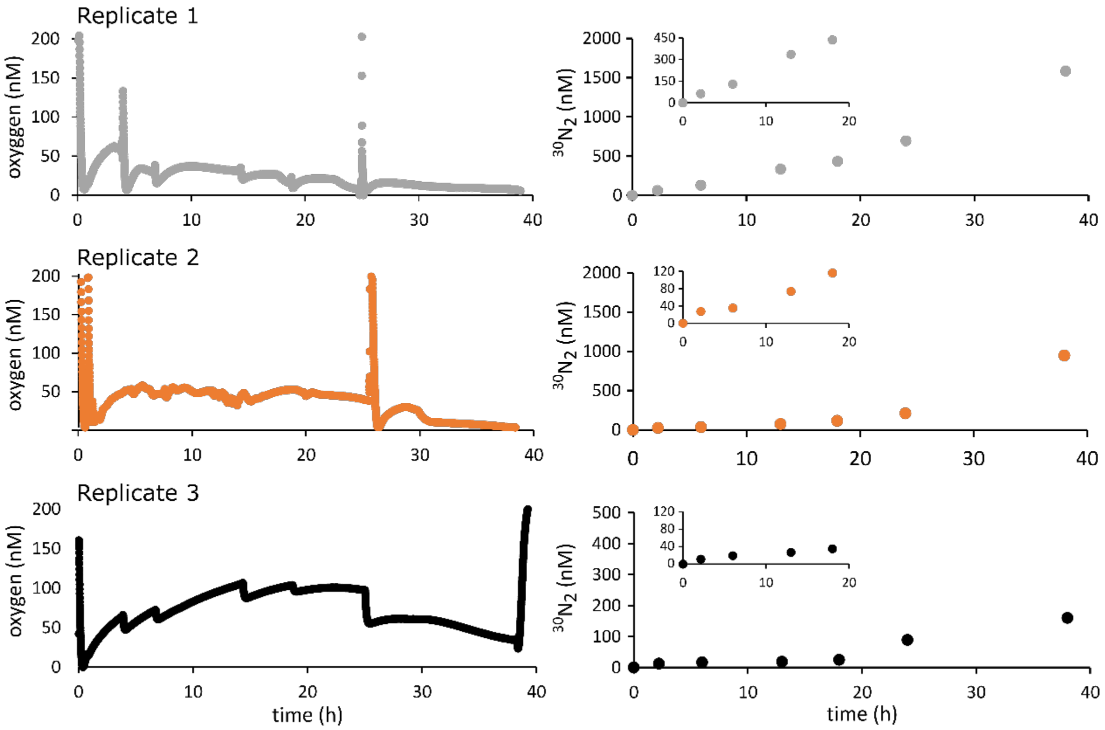


Fig. S13. Oxygen accumulation (left) and $^{30}N_2$ production from $^{15}$N-nitrite (right) in the single replicates of the incubation series 2) in fig. 3 (black diamonds). The inserts show the $^{30}N_2$ production in the first 20h of the incubation. Disturbances in the oxygen time series at 0, 2, 6, 13, 18, 24 and 38h correspond to the time points when samples for $N_2$ analysis were taken which led to slight oxygen intrusion. Only $^{30}N_2$ production is shown as no production of $^{29}N_2$ was detected (see fig 3).

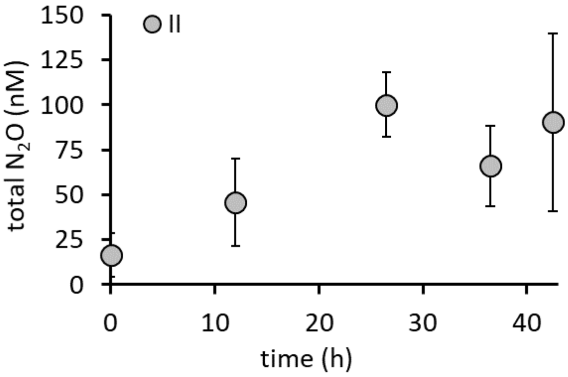


Fig. S14: Transient $N_2O$ accumulation during oxygen production in incubations of *N. maritimus.* $N_2O$ accumulation in incubation II (fig. 2) supplied with $^{15}$N-ammonium and 25 µM $^{14}$N-nitrite is shown. Due to the ongoing oxidation of $^{15}$N-ammonium to $^{15}$N-nitrite (see fig. 2) and a relatively small $^{14}$N-nitrite pool at the start, the isotopic composition of the nitrite pool changes during the incubation leading to the accumulation of $^{44}N_2O$, $^{45}N_2O$ and $^{46}N_2O$. Here, total $N_2O$ accumulation is presented. Error bars represent the standard deviation of 3 replicates.

### Supplementary Data

Data S1: Protein abundances ratios of the 40 proteins with significant increase or decrease in abundance during oxygen production compared to aerobic ammonia oxidation.

Data S2: Proteome of *N. maritimus* while aerobically oxidizing ammonia and while producing oxygen.
